# Assembling metagenomes, one community at a time

**DOI:** 10.1101/120154

**Authors:** Andries J. van der Walt, Marc W. Van Goethem, Jean-Baptiste Ramond, Thulani P. Makhalanyane, Oleg Reva, Don A. Cowan

## Abstract

**Background:** Metagenomics allows unprecedented access to uncultured environmental microorganisms. The analysis of metagenomic sequences facilitates gene prediction and annotation, and enables the assembly of draft genomes, including uncultured members of a community. However, while several platforms have been developed for this critical step, there is currently no clear framework for the assembly of metagenomic sequence data.

**Results:** To assist with selection of an appropriate metagenome assembler we evaluated the capabilities of nine prominent assembly tools on nine publicly-available environmental metagenomes, as well as three simulated datasets. Overall, we found that SPAdes provided the largest contigs and highest *N50* values across 6 of the 9 environmental datasets, followed by MEGAHIT and metaSPAdes. MEGAHIT emerged as a computationally inexpensive alternative to SPAdes, assembling the most complex dataset using less than 500 GB of RAM and within 10 hours.

**Conclusions:** We found that assembler choice ultimately depends on the scientific question, the available resources and the bioinformatic competence of the researcher. We provide a concise workflow for the selection of the best assembly tool.

## Background

The ‘science’ of metagenomics has greatly accelerated the study of uncultured microorganisms in their natural environments, providing unparalleled insights into microbial community composition and putative functionality [1]. Even though shotgun metagenomic sequencing provides comprehensive access to microbial genomic material, many of the encoded functional genes are substantially longer (∼1000 bp [2]) than the length of reads provided by the sequencing platforms [3] most commonly used for shotgun metagenomic studies (Illumina HiSeq 3000, 2 × 150 bp; http://www.illumina.com/). Thus, raw sequence data alone are typically not sufficient for an in-depth analysis of a communities functional gene repertoire. Moreover, unassembled metagenomic sequence data are fragmented, noisy, error prone and contain uneven sequencing depths [4].

To assist in the accurate and thorough analysis of metagenomes, sequence data can be assembled into larger contiguous segments (contigs) [5]. To this end, numerous metagenome assembly tools (assemblers) have been developed, the vast majority of which assemble sequences in *de novo* fashion. In short, metagenomic sequences are split into predefined segments (*k-*mers), which are overlapped into a network, and paths are traversed iteratively to create longer contigs [6]. *De novo* assembly is advantageous as it allows for more confident gene prediction than is attainable from unassembled data [7]. Furthermore, *de novo* assembled metagenomes facilitate the discovery and reconstruction of novel genomes and/or genomic elements [8].

Improvements to assembly quality have greatly expanded the scope of questions that can be answered using shotgun metagenome sequencing including, for example: determination of microbial community composition and functional capacity [9], microbial population properties [10], comparisons of microbial communities from various environments [11], extraction of full genomes from metagenomes [5] and genomics-informed microorganism isolation [12]. Each of these questions require researchers to emphasise specific features of the metagenome. Genome-centric questions [5, 12] require long contigs/scaffolds, while gene-centric questions [9–11] require high confidence contigs and the assembly of a large proportion of the metagenomic dataset.

Considering the wealth of available assemblers, it is particularly important that researchers understand assembler performance, especially for investigators who lack appropriate bioinformatic expertise. Firstly, an assembler needs to produce a high proportion of long contigs (>1000 bp). Long contigs allow for more accurate interpretation of full genes within a genomic context and facilitate the reconstruction of single genomes. A good assembler should also utilize most of the raw sequence data to generate the largest assembly span possible. Furthermore, an assembler needs an intuitive and user-friendly interface to enable assembly with minimal effort and rapid processing of the metagenomic data. Finally, tools should be able to assemble metagenomes using the least computational resources possible. Metagenomic assemblers are consistently being developed, this requires regular benchmarking, as with other bioinformatic tools [13].

Here we benchmark eight prominent open-source metagenome assemblers (Velvet v1.2.10 [14], MetaVelvet v1.2.02 [15], SPAdes v3.9.0 [16], metaSPAdes v3.9.0 [17], Ray Meta v2.3.1 [18], IDBA-UD v1.1.1 [19], MEGAHIT v1.0.6 [20] and Omega v1.4 as well as the commercially-available CLC Genomics Workbench v8.5.1 (QIAGEN Bioinformatics; https://www.qiagenbioinformatics.com/products/clc-genomics-workbench/; Supplementary Table 1). We compare each assemblers performance on nine complex metagenomes from three distinct environments (i.e., three publicly available metagenomes each from soil, aquatic and human gut niches) as well as three simulated datasets. While most of the assemblers assessed here have been tested and reviewed extensively [21–25], in this article we provide an elegant reference framework which both experienced and inexperienced researchers can use to determine which assembler is best aligned with their project scope, resources and computational background.

## Methods

### Metagenomic datasets

In this study we contrast the assemblies of nine publicly available metagenomic datasets uploaded to the MG-RAST server (http://metagenomics.anl.gov/), or the sequence read archive (SRA) (https://www.ncbi.nlm.nih.gov/sra). The metagenomes are from three distinct environments, namely; soil (Iowa [8], Oklahoma [26], and Permafrost [27]); aquatic (Kolkata Lake (unpublished data), Arctic Frost Flower [28] and Tara Ocean [29]) and human gut niches (Scandinavian Gut [30], European Gut [31] and Infant Gut [32]; Table 1). Each dataset was unique in its complexity and sequencing was performed at different depths. All metagenomes were sequenced using Illumina short read technology producing paired-end reads ranging from 100 to 151 bp in length. Most datasets were sequenced on the Illumina HiSeq 2000 platform, except for the Permafrost metagenome which was sequenced using an Illumina Genome Analyzer II, and the Kolkata Lake metagenome which comprised sequences generated by an Illumina MiSeq. This allowed for comparisons of each assemblers’ performance under different coverage and taxonomic diversity. We opted to exclusively evaluate metagenomes sequenced using Illumina platforms due to their popularity and applicability to metagenomic datasets [3].

**Table 1.** Assembly statistics and computational requirements for assembly of the Tara Oceans metagenome. Time required is given in seconds, minutes and hours for illustrative purposes and memory in GB of RAM required.

|  | Tara Ocean |  |  |  |  |  |  |  |  |
| --- | --- | --- | --- | --- | --- | --- | --- | --- | --- |
|  | CLC | IDBA-UD | MEGAHIT | metaSPAdes | MetaVelvet | Omega | Ray Meta | SPAdes | Velvet |
| Number of contigs ($\geq 500$ bp) | 50,716 | 163,815 | 216,938 | 185,419 | 67,161 | 15,982 | 6,128 | 220,178 | 57,816 |
| Total length | 46,069,409 | 179,686,756 | 210,621,485 | 202,770,058 | 55,972,515 | 34,861,819 | 7,277,214 | 275,920,632 | 45,425,460 |
| No. of long contigs ($\geq 1$ kbp) | 10,720 | 50,498 | 56,243 | 48,640 | 12,590 | 13,305 | 2,179 | 70,711 | 8,802 |
| No. of ultra-long contigs ($\geq 50$ kbp) | 0 | 2 | 1 | 37 | 0 | 9 | 0 | 54 | 0 |
| Largest contig | 39,748 | 101,400 | 62,649 | 141,519 | 30,177 | 102,255 | 41,443 | 197,381 | 21,980 |
| N50 | 880 | 1,166 | 982 | 1,124 | 805 | 2,691 | 1,329 | 1,415 | 749 |
| L50 | 14,113 | 38,236 | 58,246 | 39,033 | 21,544 | 2,737 | 1,345 | 39,617 | 19,631 |
| Mapping rate (%) | 38.98 | 52.24 | 55.92 | 64.03 | 4,117 | 13.64 | 8.25 | 64.46 | 48.19 |
| Time (seconds) | 3,527 | 69,782 | 10,455 | 125,862 | 2,527 | 168,213 | 16,419 | 80,039 | 2342 |
| Time (minutes) | 58.78 | 1,163.03 | 174.25 | 2,097.70 | 42.12 | 2803.55 | 273.65 | 1,333.98 | 39.03 |
| Time (hours) | 0.98 | 19.38 | 2.90 | 34.96 | 0.70 | 46.73 | 4.56 | 22.23 | 0.65 |
| Memory required (GB) | 16.23 | 42.84 | 10.58 | 66.53 | 109.37 | 30.7 | 42 | 157.75 | 109.37 |

Prior to assembling the short read metagenomes, we used Prinseq-lite v0.20.4 [33] for read quality control. We removed all reads with mean quality scores of less than 20 [*-min_qual_mean* 20], and removed all sequences contains any ambiguous bases (*N*) [*-ns_max_n* 0].

After quality filtering, we assessed the level of coverage of each metagenome using Nonpareil, a statistical program that uses read redundancy to estimate sequence coverage [34].

### Evaluation of the metagenome assemblers

Most assemblies were performed on a local server (48 Intel^®^ Xeon^®^ CPU E5-2680 v3 @ 2.50 GHz processors, 504 GB physical memory, 15 TB disk space) using 8 threads. However, SPAdes, metaSPAdes and IDBA-UD required more memory, and assembly was performed on the Lengau cluster of the Centre for High Performance Computing (CHPC) for the Iowa and Oklahoma soil datasets. SPAdes, metaSPAdes, IDBA-UD and MEGAHIT iteratively analyse *k*-mer lengths to find the optimal value, and these assemblers were allowed to optimise their own *k*-mer lengths. The other assemblers used *k*-mer values of 55 (Velvet: 51; MetaVelvet: 51; SPAdes: 33, 55, 71; metaSPAdes: 33, 55, 71; Ray Meta: 55; IDBA-UD: 20, 30, 40, 50, 60, 70, 71; MEGAHIT: 21, 41, 61, 81, 99; CLC Genomics Workbench: 55). In contrast to the above *de Bruijn* graph assemblers, Omega uses overlap-layout-consensus graphs to generate assemblies. Read pairs are first aligned, followed by read error correction, hash-table construction, overlap graph construction before generating contigs. We selected a minimum overlap length of 60. To control for *k*-mer length bias, we compared each assembler’s performance at *k*-mer lengths between 50 and 61. Quality of the generated assemblies were assessed using MetaQUAST. This tool calculates basic assembly statistics, including number of contigs above various lengths (500 bp, 1 kbp, 5 kbp and 50 kbp), assembly span above various lengths (500 bp, 1 kbp, 5 kbp and 50 kbp), *N*50 lengths and *L*50 lengths. To assess the accuracy and specificity of each assembler, the included synthetic metagenomes were assessed against their respective constitutive reference genomes in MetaQUAST.

To assess the volume of sequencing data that was used for each assembly, we mapped back the short fragment sequencing reads to the constructed metagenomes. This was performed using Bowtie 2 [35], using the sensitive setting. Time and memory (RAM) taken to complete assembly were calculated using an in-house bash script.

All tables and figures were drawn in R v3.4.0 or Microsoft Excel. Figure 1 was generated using the freely-available tool Nonpareil. Nonpareil estimates the percentage sequence coverage of metagenomes (as a fraction of 1) using either the forward or reverse sequence reads. These values are then plotted using a scatter plot function. Figure 2 was generated using the *heatmaply* package [36], and clustered using the *hclust* hierarchical clustering package in R. Values were calculated as a mean over- or under-representation relative to the average value obtained for all the assemblers assessed here. This provided ratios of over- or under-performance relative to the average assembly statistic (−1 to + 4). Figure 3 was generated using log-transformed data for each assembly statistic of relevance to ensure concise representation of the data.

**Figure 1.**
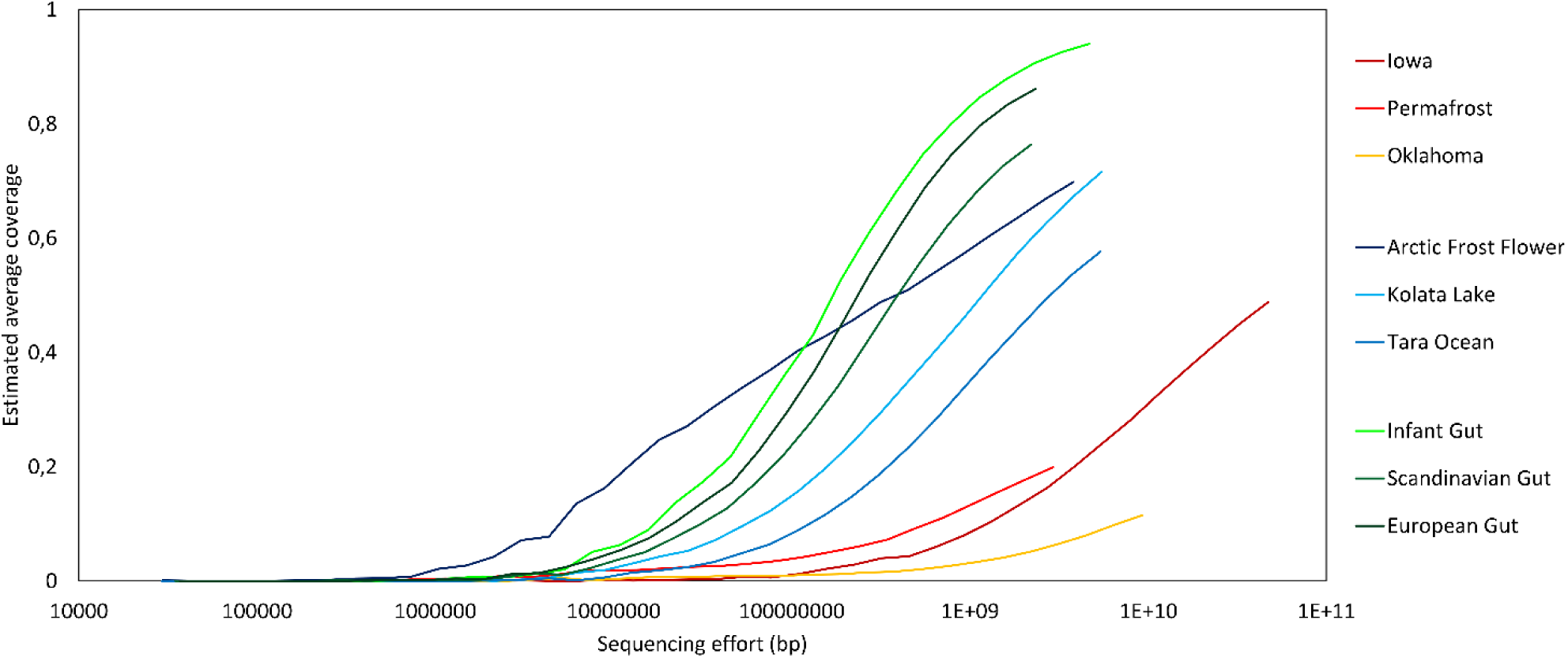
Nonpareil estimates of sequence coverage (redundancy) for the 9 metagenomes studied. Metagenomes are grouped according to their environmental niche, red colours indicate soil metagenomes, blue colours indicate aquatic metagenomes and green colours are used for human gut metagenomes. Sequencing effort is indicated in base pairs on a log scale and the estimated coverage achieved is shown as a fraction of 1.

**Figure 2.**
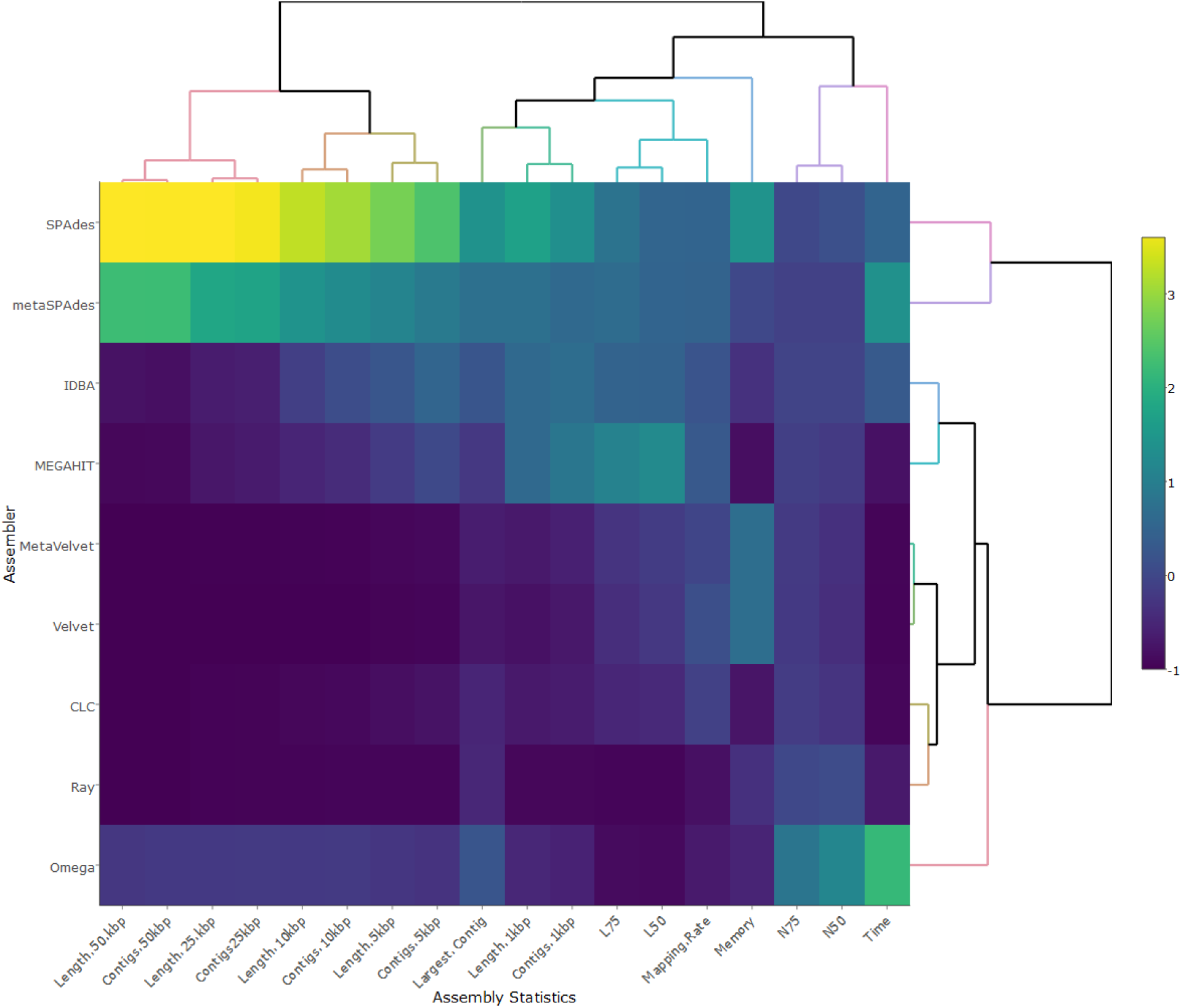
Heatmap displaying the assembly statistics measured and computational resources used by the nine tested assemblers on the Tara Ocean metagenome. Well performing statistics are shown in yellow, while dark blue regions indicate poor performance. Clustering of assemblers and assembly statistics was done using an hierarchical clustering method in R (hclust).

**Figure 3.**
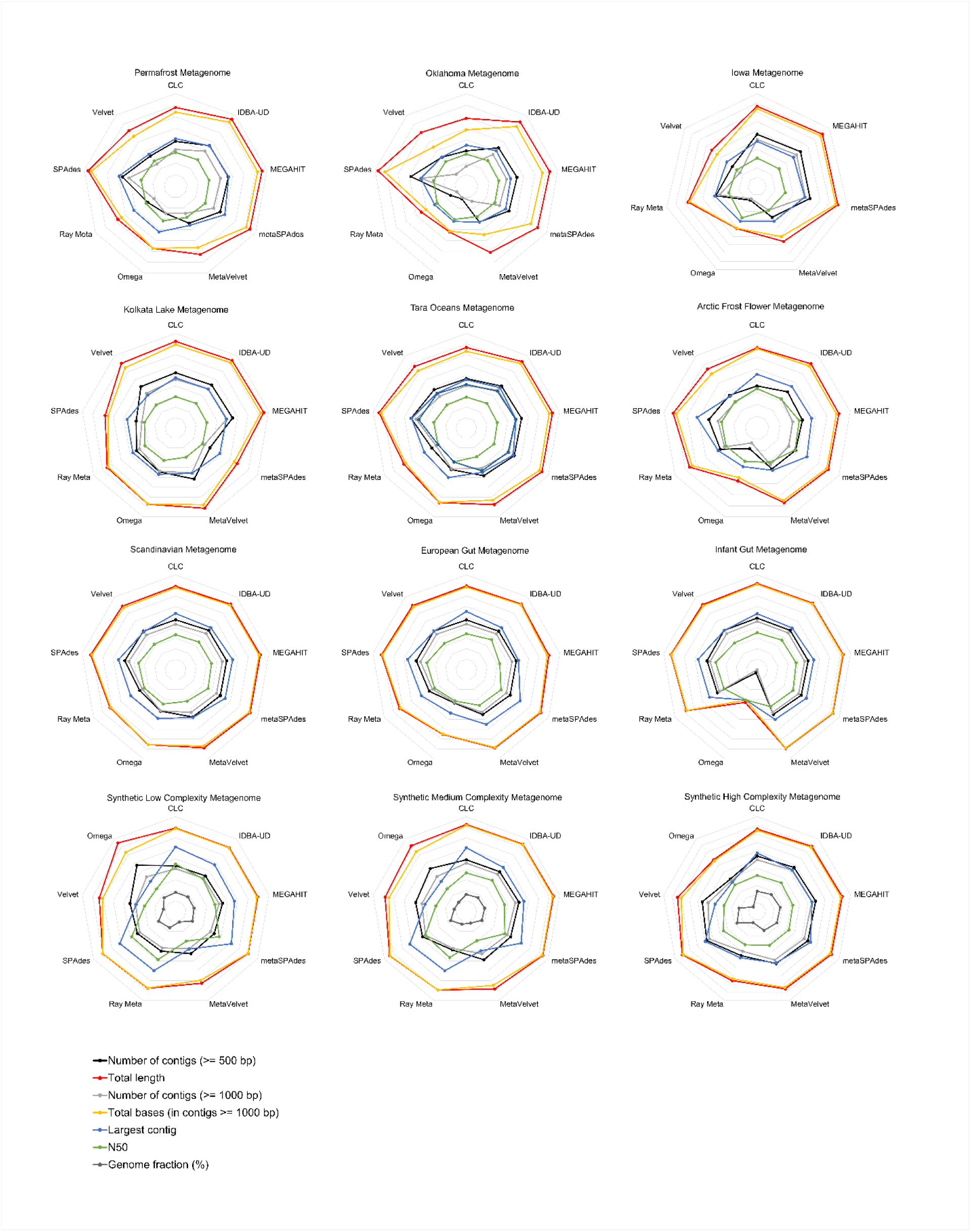
Radial plots showing assembly statistics for all metagenomes assessed as measured by the number of contigs larger than 500 bp, the total length of the assembly, the number of contigs larger than 1 kbp, the total bases calculated using only contigs larger than 1 kbp, the largest contigs, the *N50* value and for the synthetic datasets the fraction of contigs which aligned to the reference genomes provided. Metagenomes are labelled above the respective radial plots, where the first row represents the soils metagenomes, followed by aquatic, human gut and synthetic metagenomes.

Data availability is provided in Supplementary Tables 1 and 2. A link to each to assembler benchmarked is provided, as are the accession numbers for all twelve metagenomes assessed.

## Results

### Metagenome data and dataset complexity

Using Nonpareil, we confirmed that the soil metagenomes were more complex (less redundant) than the aquatic and human guts metagenomes, which were the least complex (highly redundant; Figure 1) [37–39]. All the human gut metagenomes came close to sequencing saturation (with at least 75% of the diversity sequenced; Figure 1). The infant gut metagenome was sequenced to above 90% estimated average coverage (∼94%). However, all the sequencing depths reached were insufficient to describe the complete spectrum of microbial members in the samples assessed. For example, the largest metagenome assessed here, the Iowa soil metagenome, only described 48.8% of the total microbial diversity despite the utilization of 47 Gbp of sequence data.

Estimates of the number of microbial species per gram of soil still vary substantially, with values ranging from 2000 [41] to more than 830000 [37]. These estimates do not include eukaryotic microbes, which generally possess much larger genomes and are much more difficult to fully sequence [42]. We note the published predictions that 2-5 Gbp of sequence data would fully capture an entire natural microbial community [40]. Based on our analysis, we propose that the sequencing depth required to provide comprehensive coverage of soil metagenomes should be increased by an order of magnitude, to ∼100 Gbp. This is a function of the extreme taxonomic heterogeneity of soil microbial communities, and highlights the challenge of assembling low coverage metagenomes.

### Strategy and approaches of the current research

We defined five measures to assess the performance of each metagenomic assembler tested; (1) ease of use and assembler attributes, (2) quality of assemblies generated and computational requirements, (3) influence of sequencing depth and coverage, (4) suitability to different environments and (5) their performance on metagenomes of known composition.

#### 1 Ease of use and assembler attributes

Many researchers entering the field of metagenomics are inexperienced in the use of intricate bioinformatic tools, and may lack extensive computational resources. To assess the ease of use for inexperienced computational biologists we evaluated the availability of a web application or graphical user interface (GUI), ease of installation, availability and completeness of manuals, Message Passing Interface (MPI) compatibility and programming language.

Eight of the assemblers tested here use command-line interfaces (CLI) and are open-source freeware (Velvet, MetaVelvet, SPAdes, metaSPAdes, Ray Meta, IDBA-UD, MEGAHIT and Omega). Only the commercial software CLC Genomics Workbench (Qiagen) implements a GUI (Supplementary Table 1). CLC is easily installed on most Linux, Windows or MacOS computers, whereas all other assemblers are limited to Unix-based operating systems. The GUI is intuitive, and users can assemble simply by using a point-and-click interface. CLC provides substantial support (via manuals and web based tutorials) and was the most user-friendly assembler tested here.

Unix-based assemblers are inherently more difficult to use and must be installed or compiled from source code using the CLI. All assemblers that are CLI-based can be downloaded from GitHub, while some tools (SPAdes, metaSPAdes, Ray Meta, Velvet, MetaVelvet and Omega) provide download links from their respective parent websites. All tools, barring SPAdes, metaSPAdes and IDBA-UD, provide MPI compatibility, allowing parallelization which reduces computational time. All tools assessed here provide manuals or ‘readme’ files either on their websites or GitHub repositories, although others, such as IDBA-UD, MetaVelvet and Omega, are not comprehensive and lack information on installation or implementation. Tools with more complete manuals (MEGAHIT and Ray Meta) feature extensive wiki pages and frequently asked questions. The number of citations, websites, programming languages and MPI compatibility of all the tools assessed are provided in Supplementary Table 1.

#### 2 Benchmarking quality of assemblies generated and computational requirements

Evaluating metagenome assembly quality is challenging without the use of known reference genomes for diverse microbial communities. We compared assembly quality using many standard metrics, including the total number of contigs longer than 500 bp, 1 kbp (referred to as long contigs throughout) and 50 kbp (referred to as ultra-long contigs throughout), maximum contig length, *N50* length of the contigs (length of the median contig, representing the length of the smallest contig at which half of the assembly is represented), mapping rate and assembly span (total length assembled using contigs > 500 bp). We used MetaQUAST to evaluate these assembly quality statistics [22].

We selected the Tara Ocean metagenome [29] for a comparison of each assembler at *k*-mer lengths between 50 and 61. We selected this range as the assemblers which automatically optimize *k-*mer values generally set sizes within this range. We set the other non-optimizing assemblers to 55. Compared to the other natural metagenomes, the Tara Ocean metagenomic dataset is of intermediate complexity and sequencing depth (Figure 1, Supplementary Table 2). This metagenome was sequenced on an Illumina HiSeq instrument, which is currently the most widely used shotgun metagenome sequencing technology [3]. This 5.4 Gbp metagenome comprised more than 27 million high-quality read pairs with a mean read pair length of 200.3 bp (Supplementary Table 2).

Omega (2691), SPAdes (1415), Ray Meta (1329), IDBA-UD (1166) and metaSPAdes (1124) provided assemblies with high *N50* values (> 1000 bp), while the assemblies generated using CLC, MEGAHIT, Velvet and MetaVelvet produced *N50* statistics below 1000 bp (Figure 2; Table 1). Overall, the assembly spans varied considerably with SPAdes (275.9 Mbp), MEGAHIT (210.6 Mbp), metaSPAdes (202.8 Mbp) and IDBA-UD (179.7 Mbp) assembling the largest metagenomes. Assembly span was correlated with the number of reads mapping back to the assemblies (*R*^2^ = 0.83; Supplementary Figure 3), with SPAdes and metaSPAdes having the highest values (Table 1). Both IDBA-UD and MEGAHIT mapped back more than 50% of the sequence reads to the assemblies. SPAdes also produced the most contigs over 1 kbp (70711), while MEGAHIT, IDBA-UD and metaSPAdes created fewer contigs in that size range, but all were comparable to each other (between 48640 and 56243 contigs). The largest contig was assembled by SPAdes (197 kbp), followed by metaSPAdes (142 kbp), Omega (102 kbp) and IDBA-UD (101 kbp). These three assemblers also produced the most ‘ultra-long’ contigs (> 50 kbp); with 54, 37 and 2 contigs, respectively.

The computational requirements of an assembly tool should be a major consideration when selecting an assembler. We evaluated all assemblers in relation to the time taken to assemble the Tara Ocean metagenome (Supplementary Figure 2; Table 1) using the same number of threads (*n* = 8; Supplementary Figure 2A). Velvet, MetaVelvet and CLC assembled the metagenome in less than an hour, while MEGAHIT and Ray Meta were substantially slower, assembling over multiple hours. IDBA-UD, SPAdes and metaSPAdes required considerably more time to complete assembly, taking approximately 24 hours, or more. Omega required the most time to assemble the metagenome, taking approximately 48 hours. In terms of memory requirements, SPAdes was the most ‘memory expensive’ (157 GB of RAM), followed by Velvet and MetaVelvet (both 109 GB), which is substantially more RAM than is available on an average desktop computer (16 GB). By contrast, MEGAHIT (11 GB) and CLC (16 GB) were the most memory efficient assemblers (Figure 2 and Supplementary Figure 3; Table 1).

Overall, SPAdes, metaSPAdes, IDBA-UD and MEGAHIT displayed the best performances in assembling this metagenome of intermediate size and complexity, as they produced very high *N50* values, a high proportion of long contigs and the widest assembly spans. While SPAdes was the best assembler overall, MEGAHIT was the most memory efficient, as it produced an assembly comparable to the best performing assemblers while using only a fraction of computational resources.

#### 3 Benchmarking influence of sequencing depth and coverage

Temperate soil communities are generally more diverse than extreme counterparts (e.g., permafrost; Supplementary Table 3, Figure 1) [11]. Subsequently, high levels of diversity within these biomes require much deeper sequencing effort. Differences in microorganism abundances and strain level heterogeneity introduce complications during metagenome assembly, resulting in increased memory requirements and longer computational run-times, which may challenge assemblers. The two temperate soil metagenomes assessed here have vastly different sequencing depths, thus providing us with the scope to assess the influence of sequencing depth on the performance of each assembler. The Oklahoma soil metagenome [26] had a low sequencing depth (9 Gbp) and estimated coverage (11%), c.f. the Iowa soil metagenome [8], which had a very high sequencing depth (47 Gbp) and 49% estimated coverage (Figure 1, Supplementary Table 2). We predicted that deeper sequencing effort would be correlated with an increase in metagenome coverage [34].

All assemblers successfully assembled the Oklahoma metagenome, although SPAdes required considerably more memory (up to 1 TB RAM, Supplementary Table 3). Nevertheless, SPAdes produced the best assembly statistics for most categories (9548 long contigs and an assembly span of 54.3 Mbp; Supplementary Table 3; Figure 3). IDBA-UD and MEGAHIT used less than 500 GB of RAM and were comparable in performance (3828 and 3416 long contigs, and assembly spans of 17.2 Mbp and 20.2 Mbp, respectively; Supplementary Table 3; Figure 3). It is noteworthy that while metaSPAdes was one of the best performing assemblers for the Tara Ocean metagenome (Figure 2), it performed poorly here (Supplementary Table 3; Figure 3), suggesting that metaSPAdes is ill-suited to assembling low coverage metagenomes.

The massive Iowa soil metagenome could not be assembled by either SPAdes or IDBA-UD using our available computing resources (1 TB of RAM). This is in agreement with the methodology described by the authors who generated this dataset, who digitally normalized and partitioned the Iowa metagenome to allow for assembly using Velvet [8]. Remarkably, MEGAHIT and CLC assembled the Iowa metagenome using less than 500 GB of RAM. MEGAHIT performed best across most categories tested (assembly span of 1036.5 Mbp, largest contig of 104841 bp, and 277623 long contigs; Figure 3), while CLC produced the third-best assembly (assembly span of 432.7 Mbp, largest contig of 70207 and 114196 long contigs), using less than 64GB of memory. MetaSPAdes performed comparably to MEGAHIT but had much higher computational resource requirements to assembly the Iowa soil metagenome, using up to 1TB of RAM (assembly span of 873.8 Mbp, largest contig of 188499 bp, and 225046 long contigs).

Overall, we found that sequencing depth greatly influenced the performance of the assemblers, although the most memory-efficient tools, MEGAHIT and CLC, performed well irrespective of sequencing coverage. SPAdes and IDBA-UD produced good assemblies for the Oklahoma soil metagenome, but were extremely expensive in terms of memory and failed to assemble the Iowa soil metagenome. We found that metaSPAdes produced a better assembly for the Iowa soil metagenome than the Oklahoma dataset. MetaSPAdes performed optimally for the assembly of the high-coverage metagenome, but was less efficient in the assembly of the low-coverage metagenome.

#### 4 Benchmarking suitability to various environments

Environmental samples are widely dissimilar in microbial community complexity and have distinct taxonomic compositions. In this study, we assembled metagenomes from three environmental biomes of different phylotypic complexities. Overall, SPAdes, MEGAHIT, IDBA-UD and metaSPAdes assembled most of the metagenomes well, according to the parameters we evaluated (Supplementary Tables 3-5). SPAdes consistently provided the largest contigs and the widest assembly spans. MEGAHIT demanded far fewer computational resources, and yet produced similar assemblies to metaSPAdes and IDBA-UD. CLC provided assemblies of moderate to high quality, was the easiest to use and performed particularly well on large metagenomes. Together, these results indicate that no single assembler performs best across all sequencing platforms and datasets.

#### 5 Benchmarking on synthetic metagenomes

As previously indicated, assessing metagenome assembler performance is complicated due to the unknown composition of environmental microbial communities. To overcome this challenge, we included three synthetic metagenomes of known composition to assess the error rates (such as number of indels, misassemblies, and ambiguous bases) generated by each assembler. These three metagenomes represented three discreet complexities (low, medium and high; Supplementary Figure 1), in order to challenge the assemblers with the unique properties of each dataset.

Our analysis show that more complex metagenomes led to higher error rates in the resultant assemblies (Figure 4). Notably, SPAdes produced the most misassemblies (643, 4928 and 77264 for the assemblies of low, medium and high complexity synthetic metagenomes, respectively) and the highest unaligned lengths (46 kbp, 891 kbp and 19 Mbp, respectively). IDBA-UD produced a high number of misassemblies while Omega consistently produced the most mismatches in all synthetic datasets (more than 1500 mismatches per 100 kbp for all synthetic metagenomes). CLC and Ray Meta consistently produced more than 100 ambiguous bases (N’s) per 100 kbp in each of the generated synthetic assemblies. Finally, CLC also incorporated the most indels per 100 kbp in all complexity classes (more than double the number of indels produced by any other assembler).

**Figure 4.**
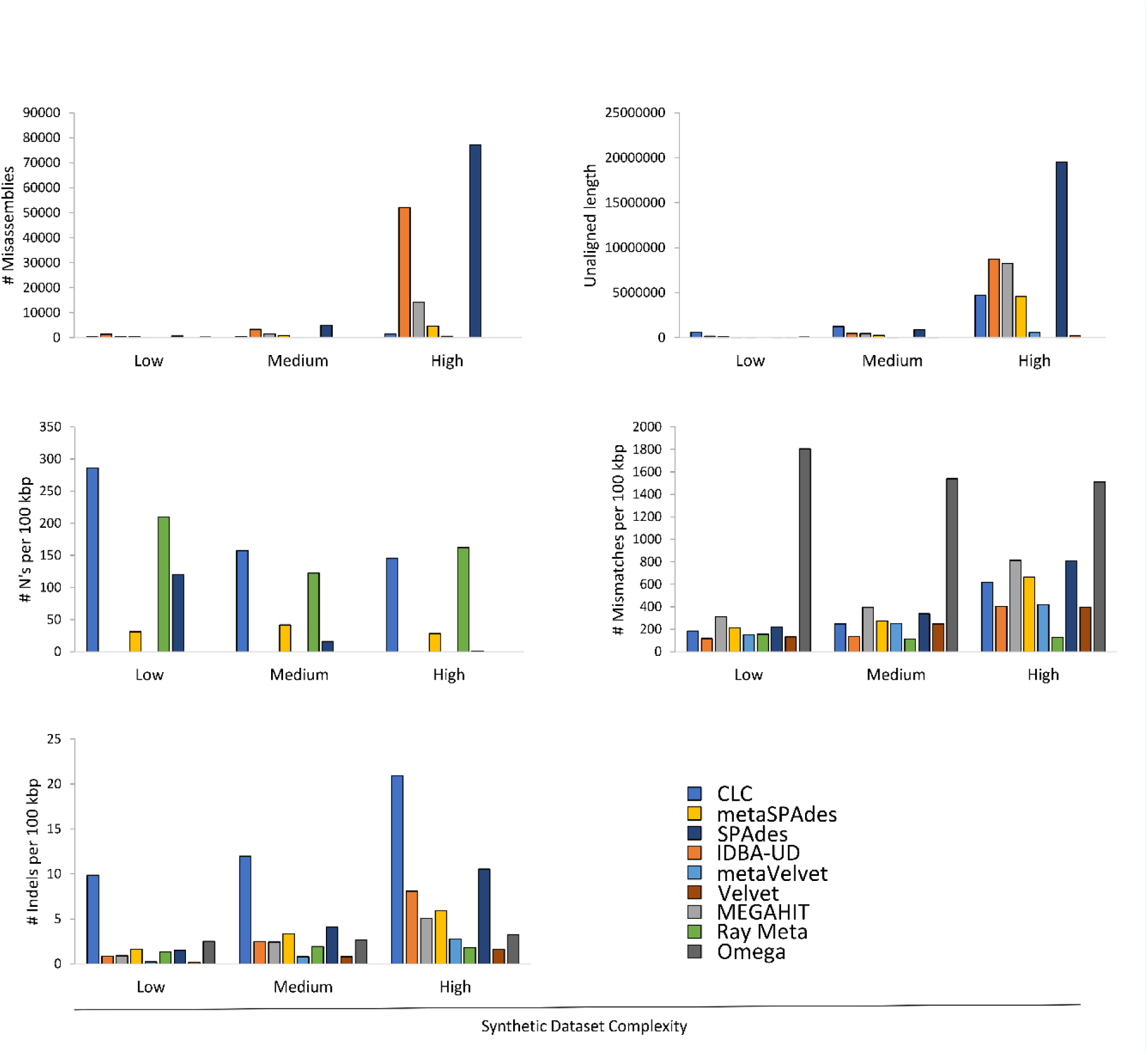
Assembler performance on synthetics simulated datasets, measured by (a) number of misassemblies, (b) unaligned length, (c) number of unassigned bases (N’s) per 100 kbp, (d) number of mismatches per 100 kbp and the number of indels per 100 kbp. These statistics represent negative assembly statistics and are a reflection of poor performance. Each assembler is indicated by different colors and the complexity of the synthetic dataset is indicated on the x-axis.

### How to select a metagenome assembler

Bioinformatics projects can be limited by memory (RAM) requirements. SPAdes, metaSPAdes, IDBA-UD, Velvet and MetaVelvet all have large memory requirements during the assembly of massive datasets. MEGAHIT, Omega and CLC are extremely memory efficient, as they required less than 500 GB of RAM to assemble the massive Iowa soil metagenome. MEGAHIT, for example, generates succinct *de Bruijn* graphs to achieve efficient memory usage [20].

Our results indicate that although many assemblers perform comparably, their applicability is defined by the research question at hand. SPAdes, for example, generated good assemblies with the most long and ultra-long contigs for most datasets. These are ideal characteristics for genome-centric studies, which require the binning of draft genomes from community sequence data [43]. By contrast, metaSPAdes considers read coverage during assembly, making it more applicable for microbial community profiling [17]. While SPAdes and metaSPAdes produced the best assemblies in general, MEGAHIT performed comparably and emerged as a rapid and memory efficient alternative assembler.

However, it should be noted that SPAdes and IDBA-UD generate high numbers of misassemblies and contigs that do not align to the reference genomes. Other assemblers such as Omega, CLC and Ray Meta each have unique error profiles, which should be considered in light of the research questions asked. For example, when assessing strain level genomic variations (SNP’s), assemblers that generate high numbers of indels and mismatches should be avoided. In addition, while SPAdes generates many mismatches, if the aim is to extract single genomes from a metagenome, manual curation of the newly re-constructed draft genomes will identify and correct such misassemblies.

In conclusion, we argue that when selecting an assembler, the primary consideration should be the research question. Selecting an appropriate assembler is essential to make full use of metagenomic sequence dataset. The primary objectives of the project, whether gene- or genome-centric, for example, should dictate the choice of assembler. We suggest that a secondary consideration should be the computational resources available to the researcher. Some assemblers are very memory efficient, while others sacrifice computational resources for improved assembly quality. Finally, as most assemblers use a CLI (and are more flexible than those constrained by a GUI), the GUI-based CLC platform is an excellent alternative if bioinformatic skill level is a consideration.

## Other analyses

In additional analyses (Figure 3), we compared the performance of each assembler on a low diversity soil metagenome (Supplementary Table 3), other aquatic metagenomes (Supplementary Table 4) and human gut microbiomes (Supplementary Table 5).

## Discussion

Over the last decade, high throughput sequencing has revolutionised the field of microbial ecology [44]. Amplicon-based technologies have allowed for near-complete classification of whole microbial communities, including populations of bacteria, archaea and fungi [45]. The emergence of two key platforms for analysing amplicon sequencing data, mothur [46] and QIIME [47], has allowed for methodological standards to be set, which enables robust comparisons between studies [48].

While whole community shotgun metagenome sequencing has facilitated the in-depth description of microbial communities from diverse environments, such as the human gut [49] and acid mine drainage systems [50], no standards exist with regard to assembly platforms or their use. While numerous reviews on strategies to analyse metagenomic data have been published [51], there are currently no standard assembly procedures implemented to enable thorough comparative analyses between projects. Numerous pipelines for processing metagenomic sequence data are available. These typically integrate existing tools into a single workflow for rapid, standardized analysis (e.g., MG-RAST, MetAMOS, and IMG/M) [52–54]. However, few of these pipelines are as widely used as mothur or QIIME in barcoding studies. This is partly because integrated metagenome analysis tools, such as MetAMOS, do not achieve the flexibility afforded by using each tool individually (e.g., using separate tools for assembly, binning and taxonomic assignment).

Consequently, investigators can analyse unassembled reads [11], optimize their assembly parameters or even develop their own tools to assemble their data prior to further analysis [55]. However, within the scope of metagenome assembly, essential details are often omitted when describing methods [56]. This leads to methodological discrepancies, and severely limits the possibility of making routine, robust comparisons between studies. This issue was recently highlighted by J Vollmers, S Wiegand and A-K Kaster [21] and WW Greenwald, N Klitgord, V Seguritan, S Yooseph, JC Venter, C Garner, KE Nelson and W Li [57] who reported that the taxonomic diversity patterns of microbial communities differed substantially, depending on the assembler used. While some recent studies have applied single cell sequencing [58] or chromosome capture [59] approaches to enhance metagenome assembly, these techniques remain inaccessible to most researchers. We provide an evaluation of commonly-used assemblers on standard shotgun sequenced metagenomes.

In our comparative analyses of the most popular assembly platforms, SPAdes produced the most long contigs, independent of the metagenome origin. However, this assembler introduced a large number of misassemblies in high complexity datasets. SPAdes is ideal for genome-centric research questions that require long and ultra-long contigs, such as those that aim to bin and reconstruct single genomes from shotgun metagenomes [16]. By contrast, MEGAHIT and metaSPAdes provided very large assembly spans and consider sequence coverage during assembly, reducing the number of misassemblies generated. IDBA-UD also produced large assembly spans and a high number of contigs, but at the cost of generating misassemblies for complex datasets. These tools are thus more appropriate for research questions related to taxonomic profiling of natural microbial communities, for functionally annotating microbial communities, for the analysis of population scale dynamics or for comparison of microbial communities across biomes [17, 20]. By analysing metagenomes of known composition and complexity, we found that each assembler tested here generated a unique error profile (e.g., IDBA-UD produces many misassemblies, CLC produces many indels and Omega produces many mismatches). As mentioned above, this excluded some assemblers from specific research objectives (e.g., using CLC for variant calling). This reiterates the fact that the research question should be the primary consideration when selecting the appropriate assembler, and that these assembler-specific drawbacks should also be considered.

Overall, MEGAHIT produced some of the best assemblies throughout this study, while only using a fraction of the computational resources required by other assemblers. We strongly recommend MEGAHIT for researchers who do not have access to large computational resources. Finally, the CLC assembler is ideal for researchers who lack a depth of bioinformatic knowledge, or who prefer to use a GUI and are willing to invest in software which is easier to use. CLC is easy to install, has an intuitive interface and provides a compromise in which assembly quality may be sacrificed for ease of use. Strikingly, the most widely cited assembler assessed here (Velvet cited 5974 times; Supplementary Table 1) did not perform well across most metagenomes, while scarcely cited platforms (MEGAHIT, metaSPAdes cited 114 and 18 times, respectively; Supplementary Table 1) performed well across most statistics assessed here.

## Conclusions

No assembler tested here consistently provided superior assemblies across the different metagenomes. Consequently, we propose a viable methodology for the selection of an appropriate assembler, dictated by (1) the scientific research question posed, then by (2) the computational resources available, and (3) the bioinformatics skill level of the researcher (Figure 5). In light of the above proposed framework, we urge researchers to carefully consider the assembler used (as well as the entire bioinformatics pipeline followed) while specifically bearing in mind their research question and what feature of the dataset they want accentuated.

**Figure 5.**
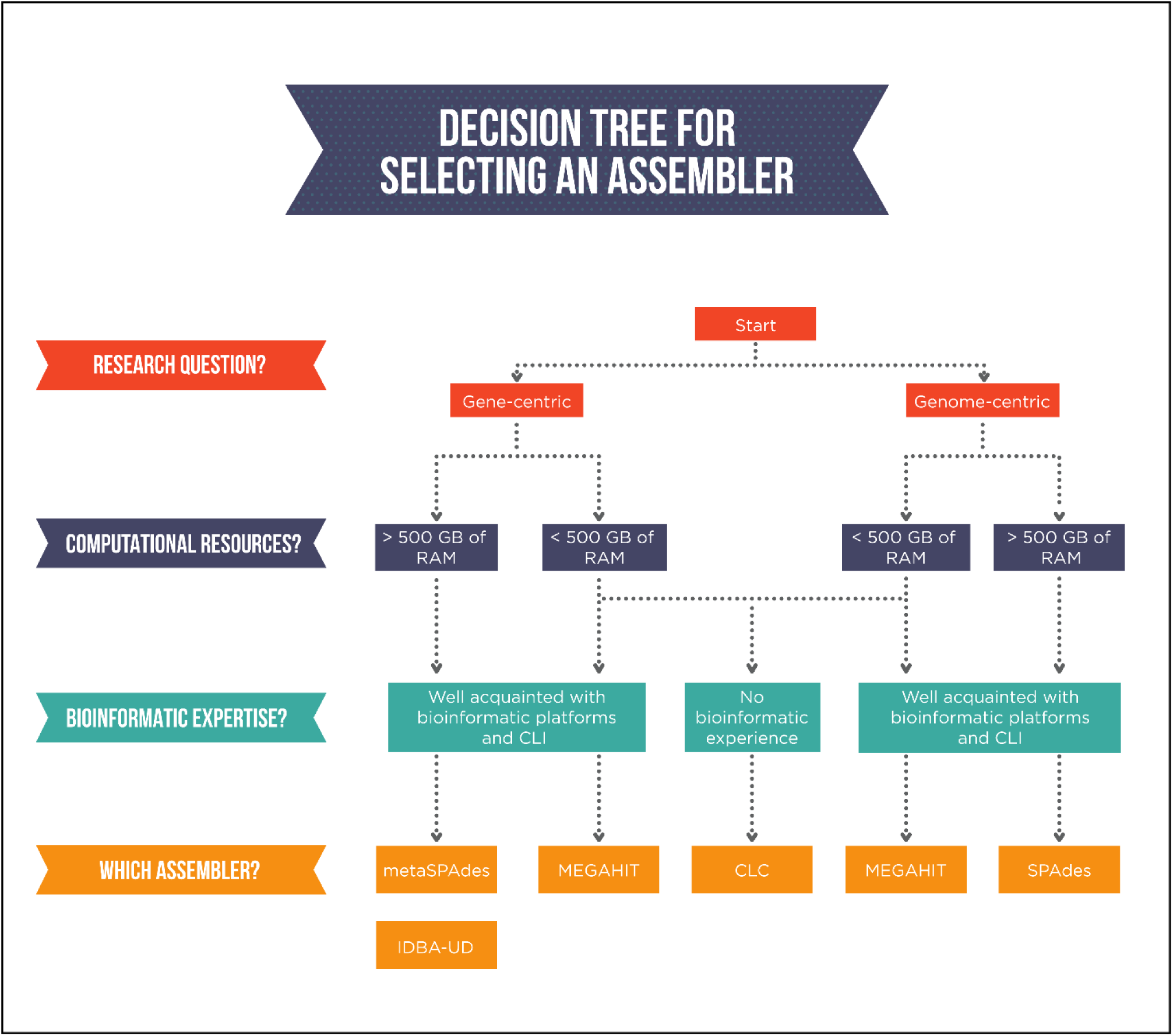
Proposed workflow to select a metagenome assembler based on the research question, the computational resources available and the bioinformatic expertise of the researcher.

## List of abbreviations

contigs: contiguous segments
SRA: sequence read archive
MG-RAST: Metagenomics Rapid Annotation Server
CHPC: Centre for High Performance Computing RAM random access memory
GUI: graphical user interface
MPI: Message Passing Interface
CLI: command-line interface
SNP: single nucleotide polymorphism

## Declarations

### Ethics approval and consent to participate

Not applicable

### Consent for publication

Not applicable

### Availability of data and material

The datasets analysed during the current study are available in the Sequence Read Archive (SRA) repository, under accession numbers SRR351474, SRR958082, ERR598950, ERR526087, SRR341725 and ERR732914. As well as the MG-RAST repository, under accession numbers 4470009.3 - 4470010.3, 4673644.3 - 4673645.3 and 4537104.3 - 4537105.3. Synthetic simulated metagenomes are available from https://data.cami-challenge.org.

### Authors’ contributions

AJvdW contributed most to research, analysis, formulation of ideas and writing. MWVG contributed research, analysis, formulation of ideas and writing. JBR contributed formulation of ideas, direction and writing. TPM contributed formulation of ideas and writing. OR contributed direction and writing. DAC contributed formulation of ideas, direction and writing.

## Acknowledgements

We thank Dr Surendra Vikram for helping with the installation of assemblers, as well as Dane Kennedy from the CHPC, South Africa with his assistance in accessing the large memory nodes of the CHPC. We gratefully acknowledge support from Ms Amanda van der Walt help with improving the quality of our figures. Finally, we thank S. de Scally and A. E. Visser for their support and helpful discussions.

## Additional Files

### Supplementary Tables and Figures

**.pdf**

**Title of data:**

**Supplementary Table 1. Attributes of *de novo* assemblers used in this study**. Included in this table are the versions of each assembler used in this study, along with the release date of each version. We provide a link to each assemblers’ website accompanied by its reference and number of citations. We gauge ease of use by providing the programming language and MPI compatibility of each tool as well as assessing the completeness of each tools’ available documentation.

**Supplementary Table 2. Characteristics of the metagenomic datasets used in this study.** Three metagenomes from three distinct environments (Soil, Aquatic and Human gut) were selected, and we provide accession numbers, sequencing platforms used and basic sequence characteristics (pre- and post-filtering) of each metagenome.

**Supplementary Table 3. Assembly statistics for the assembled aquatic metagenomes.**

**Supplementary Table 4. Assembly statistics for the assembled soil metagenomes.**

**Supplementary Table 5. Assembly statistics for the assembled human gut metagenomes.**

**Supplementary Table 6. Assembly statistics for the synthetic metagenomes.**

**Supplementary Figure 1. Nonpareil estimates of sequence coverage (redundancy) for the 3 synthetic metagenomes studied.**

**Supplementary Figure 2.** Computational requirements for the Tara Ocean metagenome. A) Total assembly span proportional to wall time required. B) Total assembly span in relation to peak memory usage.

**Supplementary Figure 3.** Correlation between assembly span and mapping rate. The exponential trendline indicates a very strong positive correlation between the amount of data utilized and the size of the generated assembly (R^2^ = 0.83).

